# Developmental changes in the processing of faces as revealed by EEG decoding

**DOI:** 10.1101/557447

**Authors:** Inês Mares, Louise Ewing, Emily K. Farran, Fraser W Smith, Marie L Smith

## Abstract

Rapidly and accurately processing information from faces is a critical human function that is known to improve with developmental age. Understanding the underlying drivers of this improvement remains a contentious question, with debate continuing as to the presence of early vs. late maturation of face-processing mechanisms. Recent behavioural evidence suggests an important ‘hallmark’ of expert face processing – the face inversion effect – is present in very young children, yet neural support for this remains unclear. To address this, we conducted a detailed investigation of the neural dynamics of face-selective processing in children spanning a range of ages (6 – 11 years) and adults. Uniquely, we applied multivariate pattern analysis (MVPA) to the electroencephalogram signal (EEG) to test for the presence of a distinct neural profile associated with canonical upright faces when compared both to other objects (houses) and to inverted faces. Results revealed robust discrimination profiles, at the individual level, of differentiated neural activity associated with broad face categorization and further with its expert processing, as indexed by the face inversion effect, from the youngest ages tested. This result is consistent with an early functional maturation of broad face processing mechanisms. Yet, clear quantitative differences between the response profile of children and adults is suggestive of age-related refinement of this system with developing face and general expertise. Standard ERP analysis also provides some support for qualitative differences in the neural response to inverted faces in children in contrast to adults. This neural profile is in line with recent behavioural studies that have reported impressively expert early face abilities during childhood, while also providing novel evidence of the ongoing neural specialisation between child and adulthood.

## 1. Introduction

Human faces provide a wealth of social information that powerfully informs our behaviour. Our sensitivity to these cues starts emerging very early in life; a remarkable preference for selectively attending to face-like visual stimuli has been reported in newborns (Johnson, Dziurawiec, Ellis, & Morton, 1991) and more recently even in foetuses (Reid et al., 2017). Unsurprisingly, these early perceptual biases do not match the sophistication of face abilities observed later in development. Studies tracking outcomes on lab-based face processing tests in the early years of life report improvements in performance with age (e.g., Carey, Diamond, & Woods, 1980; Hills & Lewis, 2018; Laurence & Mondloch, 2016; Lawrence et al., 2008; Mondloch, Le Grand, & Maurer, 2002), peaking at around 30 years (Germine, Duchaine, & Nakayama, 2011). Fierce debate continues, however, regarding the mechanism/s driving the observed change (see McKone, Crookes, Jeffery, & Dilks, 2012 for an extensive review).

There are two contrasting perspectives on this issue. One hypothesis suggesting late maturation of face-selective abilities proposes that domain-specific mechanisms undergo tuning with face experience, leading to progressively more sophisticated face processing capacity with increasing age (e.g. Carey & Diamond, 1977; Germine et al., 2011; Hills & Lewis, 2018; Susilo, Germine, & Duchaine, 2013). In contrast, a hypothesis of early face-selective maturation contends that any observed changes in performance during development reflects maturation of general cognitive processes that are not face-selective (Crookes & McKone, 2009; McKone et al., 2012), e.g., improvements in children’s attention, memory and executive functioning across childhood are well-documented (Casey, Giedd, & Thomas, 2000; Zelazo & Mller, 2002).

Early empirical evidence tended to support the former, a late maturation of face specific abilities. For example, disproportionate performance costs are associated with the inversion of faces, compared to other objects, in adults (e.g. Yin, 1969). This face inversion effect has been taken to reflect, in part, specialised holistic processing for upright faces (Edmonds & Lewis, 2007; Farah, Tanaka, & Drain, 1995; Freire, Lee, & Symons, 2000; Maurer, Grand, & Mondloch, 2002). Relatively attenuated or absent face inversion effects in young children appear to suggest an initially immature holistic processing of faces that is reliant on a non-expert processing strategy for faces at both orientations (Carey & Diamond, 1977; Hills & Lewis, 2018; Schwarzer, 2000)(Carey & Diamond, 1977)(Carey & Diamond, 1977)(Carey & Diamond, 1977)(Carey & Diamond, 1977)(Carey & Diamond, 1977)(Carey & Diamond, 1977). In particular, researchers have suggested that children rely to a greater extent on individual facial features than adults, who employ a more holistic processing strategy for upright faces (see Carey & Diamond, 1977; Carey et al., 1980).

Contemporary research has, however, begun to challenge this notion of qualitative differences in the face processing of children and adults. In particular, researchers have highlighted methodological limitations in these earlier studies, e.g., failure to adequately match task difficulty for adults and young children (e.g., see Crookes & McKone, 2009; McKone et al., 2012). Taking these concerns into account, more recent developmental studies suggest that the magnitude of the face inversion effect is in fact similar between childhood (7 years of age or earlier) and adulthood (Crookes & McKone, 2009; McKone et al., 2012). Converging evidence from contemporary infant research also indicates that this marker of specialised face processing may be present from 1 to 3 days after birth, with infants showing susceptibility to two tests of holistic face processing: the Thatcher illusion (Leo & Simion, 2009) and the composite effect (Turati, Di Giorgio, Bardi, & Simion, 2010). Taken together, these results suggest that this key hallmark of expert face processing may be present, at least *qualitatively*, in infancy and early childhood, supporting an early maturation of face specific abilities.

Typically used behavioural measures, such as reaction time and accuracy, reflect the summation of children’s cognitive, perceptual and motor processes. Clear interpretation of performance differences on such measures are therefore complicated by the possibility of different rates of maturation across these distinct processes. Investigating the neural markers associated with the development of face-processing should bypass these issues and provide explicit evidence confirming the presence (or absence) of neural indicators of face specific abilities.

Indeed, both Functional Magnetic Resonance Imaging (fMRI) and electroencephalography (EEG) results support face-selective neural development during childhood, i.e. alterations in face-related neural activity. Despite methodological concerns (e.g. the use of adult size head coils, see McKone et al., 2012), fMRI studies consistently observe increases in the size and face-selectivity of key neural regions (e.g., the fusiform face area) with increasing age (e.g., Golarai et al., 2007; Passarotti, Smith, DeLano, & Huang, 2007; Scherf, Behrmann, Humphreys, & Luna, 2007). Further some electroencephalography (EEG) evidence in young infants does indicate specialised cortical processing of upright human faces, compared to inverted faces, noise, or faces of other species (e.g., monkeys) from the first year of life (Halit, Csibra, Volein, & Johnson, 2004; Halit, de Haan, & Johnson, 2003). However, relatively little EEG research has investigated developmental changes during childhood in the time course of face processing (Itier & Taylor, 2004b, 2004a; Kuefner, de Heering, Jacques, Palmero-Soler, & Rossion, 2010; Miki, Honda, Takeshima, Watanabe, & Kakigi, 2015; Taylor, Edmonds, McCarthy, & Allison, 2001; Taylor, McCarthy, Saliba, & Degiovanni, 1999) and the results of the few studies conducted have been mixed.

Basic face categorization effects i.e., a selective neural response to faces compared to other objects is routinely observed in the typically analysed electrophysiological ‘hallmark’ of face sensitivity, the N170 component (Bentin, Allison, Puce, Perez, & McCarthy, 1996), from four years of age and show limited signs of further developmental change (Kuefner et al., 2010). By contrast, studies evaluating face inversion effects on the N170 component (i.e., a selective neural N170 response to upright compared to inverted faces, which is very robust in adults) have produced conflicting evidence. Though face-orientation sensitivity has been found in one study in children as young as 5 years of age (Melinder, Gredebäck, Westerlund, & Nelson, 2010), several others concluded that differences emerge only after 10 years (Miki et al., 2015) or report that the pattern and directionality of the face-inversion effect over the N170 component changes during development and may even disappear between the ages of 10 and 11 (Itier & Taylor, 2004b, 2004c). These highly variable neural findings stand in stark contrast to the emerging pattern of qualitatively mature behavioural face inversion effects in children from 4 to 6 years.

It is notable too that the few existing EEG studies to date have focused on a restricted subset of face-related components. Typically, this has been the N170 and the P100 component, a component originating in extrastriate visual areas (Di Russo, Martínez, Sereno, Pitzalis, & Hillyard, 2002) linked to low-level stimulus properties and attention. The P100 component has however also been shown to be face sensitive in children, with faster and larger responses to faces than other objects, and faster but smaller responses to inverted than upright faces (Kuefner et al., 2010; Taylor, Batty, & Itier, 2004b). After presentation of a test stimulus, these components are averaged from the neural activity recorded from a small, targeted set of electrodes, over a specific time-window. Such an approach is standard in EEG research, but is not ideal for analysing developmental changes due to particularly high temporal (Taylor, Batty, & Itier, 2004a) and spatial variability in neural activity across individual children and between age groups (Scherf et al., 2007).

Here we sought to provide a more comprehensive understanding of the neural development of face processing abilities. In addition to the traditional approach of investigating ERP face sensitive markers, we employ multivariate pattern analysis (MVPA) to characterise face-related neural representations (see, Nemrodov, Niemeier, Mok, & Nestor, 2016; Smith & Smith, 2018). MVPA has only relatively recently been applied to explore the time-course of neural representations from time-sensitive neuroimaging approaches (EEG and MEG, see Grootswagers, Wardle, & Carlson, 2017 for a review), and never before with a developing sample. Our application of MVPA in this context permits a uniquely broad analysis of face selective neural activity, freed from a priori constraints such as predetermined time windows and a small number of electrodes typically showing maximal response for the ERP components of interest (important factors in any traditional analysis). MVPA rather makes use of the overall pattern of neural activity measured from a broad set of electrodes across the cortex (e.g. all recording electrodes or a large set of electrodes covering for example visual areas) and as such is not limited by specification of electrode location. The approach is thus particularly well suited to probing the stability of face selective processing across development, where there is reported to be considerable variability in the neural sources contributing to category sensitivity (Scherf et al., 2007).

We recruited a wide age range of participants (6 to 11 years and adults) and assessed their neural responses to upright and inverted faces and houses. To avoid potentially confounding differences in cognitive ability between age groups, participants completed an orthogonal task unrelated to the faces or houses. We employed MVPA and standard ERP analysis to explore the representation of face category information (contrasting upright faces and houses), and more specific face expertise (contrasting the canonical upright face configuration and inverted faces). If the improvements widely observed on behavioural measures of face processing reflect only changes in general cognitive functioning, then we should see few specific changes in how the brain responds to these different stimuli categories across time in the absence of task demands (i.e. children’s neural response should demonstrate an adult-like pattern of differentiated neural activity for faces vs. other objects: indexing basic category sensitivity, as well as for upright faces vs. inverted faces: indexing face expertise). Alternatively, however, if face processing ability is refined with age and experience, then we should observe age-related changes in the neural selectivity to these categories across childhood - particularly so for the more experience-sensitive face inversion effect.

## 2. Methods

### 2.1. Participants

A total of 99 participants were initially recruited and tested, from across four age groups, 6 to 7 year olds (N=26), 8 to 9 year olds (N=27), 10 to 11 year olds (N=23) and adults (N=23). Of these, a subset of participants provided sufficient artefact free EEG data to consider further (defined as a minimum of 30 clean trials per experimental condition): 17 participants aged 6 to 7 years, 15 aged 8 to 9 years, 21 aged 10 to 11 years, and 19 adults. Further, to approximately equate the sample size between groups (and therefore equate the sensitivity of the approach), we matched the two older age groups (10 - 11 year olds, adults) with the number of younger children so that the final sample comprised 17 individuals at age 6-7 (9 female, mean age, 86.53 months std = 5.3, 77 to 95 months), 15 individuals aged 8-9 (9 female, mean age, 109.00 months, std = 8.66, 96 to 119 months), 17 aged 10-11 (9 female, mean age = 132.47 months, std = 7.23, 122 to 142 yrs) and 17 adults (10 female, mean age 26.4 yrs, std = 3.5, 22 to 34 yrs).^1^ Written informed consent was obtained from all adult participants as well as from the children’s parents according to the Declaration of Helsinki. This study was approved by the ethical committee of the Department of Psychological Sciences, Birkbeck College, University of London. Adult participants were compensated for their time either with course credits or a small monetary reimbursement. Child participants were awarded a ‘Junior Scientist’ certificate and surprised with a small-value book token upon completion of their experimental session.

### 2.2. Stimuli

Six unique face identities with neutral face expressions were presented (standardized greyscale photographs from Schyns & Oliva, 1999) alongside greyscale photographs of six unique houses (photographs from (Eimer, 2000), similarly edited to have the same outline as the face stimuli). Luminance and contrast were controlled for using the Shine toolbox (Willenbockel et al., 2010). Inverted versions of the upright images were created for all stimuli. Participants sat 70cm from the computer screen such that stimuli subtended around 4.09° width by 6.13° height degree of visual angle.

### 2.3. Procedure

Participants completed the EEG recording as part of a larger battery of tasks administered during a 90 – 120 minute testing session, with breaks. Participants were seated comfortably in a chair in an electrically shielded and sound-proofed room throughout the task. They were accompanied at all times by an experimenter who guided them through the task (and preparation), providing encouragement and ensuring that breaks were taken whenever required. We used Eprime software, version 2.0 (Psychology Software Tools Inc.; www.pst-net.com/eprime) to centrally present each stimulus on a grey background (750 ms) followed by a black fixation cross (for between 1700 and 1900 ms in steps of 25 ms). Participants completed 60 trials of each condition (faces and houses, upright and inverted), for a total of 240 trials. They were asked simply to view each image closely and look out for brightly coloured butterflies that appeared to the left or right of fixation on 60 additional catch trials (for a total of 300 experimental trials). To maintain interest and attention, participants made a speeded keyboard response to indicate whether these butterflies appeared on the left or right side of the screen. Participants took short breaks between each of 10 × 30-trial blocks (24 faces/houses, 6 butterflies). The experimenter also closely monitored task engagement and discontinued the experiment where there were concerns about task engagement or fatigue.^2^ We note that, as part of a larger battery of tasks, prior to participation in the main EEG experiment, participants also completed a short familiarisation task in which they were asked to learn by name three of the six used identities (see Smith et al, 2017 for details of the familiarization task - the puzzle bubble game).^3^ No information was given to participants regarding the faces shown in the main experiment e.g. to note gender or familiarity, rather they were asked to pay attention to the stimuli on the screen and only respond to the relatively rare appearance of a butterfly. While the potential effect of familiarity is certainly an interesting question, given the insufficient statistical power this orthogonal categorization was not analysed via ERPs or MVPA.

### 2.4. EEG recording and analysis

EEG was continuously recorded using a fitted cap (EASYCAP) with 32 Ag-AgCl electrodes placed according to the international 10/10 system. Electrode impedance was lowered below 10 kΩ and an additional electrode was placed below one of the eyes to monitor vertical eye movements and blinks. EEG was acquired at a sampling rate of 500 Hz, with electrode FCz acting as the reference and AFz as ground. Data was analysed using the Matlab toolbox EEGLAB (Delorme & Makeig, 2004).

Continuous data was band pass filtered between 0.1 and 40 Hz, and epoched around stimulus onset from −200 ms to 500 ms. Rejected channels due to noise were interpolated (maximum 4; M=2.33±1.34 channels). Epochs were baseline corrected using the 200 ms previous to stimulus onset. Test trial epochs (catch trials were excluded from the analysis) were visually inspected to remove artefacts (eye blinks/movements, muscle activity). After initial artefact rejection (14.12±1.18% of each participants total trials), the mean number of trials was equalized across the four age groups (218 trials)^4^ to further equate sensitivity of the subsequent analysis.

Channels for ERP analysis were selected based on the maximum peak difference between P100 and N170 from the average of all conditions over parieto-occipital channels (O1/2 and P7/8). Mean amplitude was calculated for the P100 in a 20 ms window centred around the average P100 peak for each group (6-7 yrs, 126 ms; 8-9 yrs 126 ms; 10-11 yrs 124 ms and adults 102 ms). A similar approach was conducted for the N170 component using a 40 ms window given the relative broader form of this component (6-7 yrs, 200 ms; 8-9 yrs 184 ms; 10-11 yrs 184 ms and adults 162 ms). P100 peaks were identified for latency analysis as the maximum positive peak in a window between 70 ms and 178 ms after stimuli onset. One participant aged 10-11 yrs was removed from this latency analysis due to the lack of identifiable P100 peaks in all conditions. N170 latency was not analysed due to the frequent presence of a bifid peak, as has previously been described in young children (Taylor, Batty & Itier, 2004).

### 2.5. MVPA Analysis

We used MVPA to reveal whether distinct patterns of neural activity are associated with the processing of our categories of interest. That is, we sought to determine whether a model can predict whether a participant was viewing a particular stimulus, e.g., an upright vs inverted face. If it can, then we are able to infer that the electrophysiological data contains information pertinent to the distinct representation of these two categories (see Grootswagers et al., 2017)). Linear support vector machine (SVM) classifiers were trained on single trial ERPs across all time samples (downsampled to 250Hz) using occipital electrodes (O1, O2, P7, P8, P3, P4, Pz, TP9, TP10) for each of the three planned binary comparisons (i.e. 50% chance level): upright faces vs. inverted faces; upright faces vs. upright houses; upright houses vs. inverted houses. We chose to use this set of occipito-temporal electrodes as we expected them to cover the activity generated in the key brain areas important in visual processing of our face stimuli (see also Smith & Smith, 2018).

For each classification problem (e.g. upright vs inverted), the classifier was trained and tested on independent sets of data. We used 70% of data for training and 30% for testing (see Smith & Smith, 2018) and classification performance was calculated using a twenty-fold cross-validation. This procedure was repeated 100 times for robustness (Cauchoix, Barragan-Jason, Serre, & Barbeau, 2014), effectively meaning 200-fold cross-validation. Accuracy was calculated by testing the trained classifier against the averaged EEG pattern across all trials from the test set of each respective condition, as a means of increasing signal to noise (Gallivan, McLean, Valyear, & Culham, 2013; Smith & Muckli, 2010; Smith & Smith, 2018). To produce an empirical measure of the chance level we performed the same procedure on permuted labels (100 iterations). A classifier using the true labelling was also included in the distribution of results as one of the possible outcomes. Averages were created from the 100 iterations of the classifiers created with the correct and permuted labels. Significant decoding was computed at the group level via a paired samples t-test across all participants (one-tailed) for each time point that tested whether the average observed decoding was significantly higher than the average chance level decoding (False Discovery Rate, FDR, corrected).^5^

We then sought to extend our investigation of group level category decoding of these same three comparisons at the individual participant level. To establish significant decoding at the individual level, a further 900 iterations of the classifier were generated per participant using permuted labels in order to create a null distribution per participant (total of 1000 permutations). The individual participant probability was then calculated as the proportion of the null distribution that was greater than or equal to the accuracy obtained with correct labels, with significant classification being considered when the accuracy obtained with correct labels is greater than or equal to 95% of the null distribution (FDR corrected, see Pereira, Mitchell, & Botvinick, 2009; Smith & Muckli, 2010).

At the individual level we then extracted four metrics: *decoding onset* - defined as the time-point where significant decoding first surpassed chance levels (FDR corrected) and exceeded baseline levels, *sustainability of decoding* - defined as the percentage of significant decoding in a given time-window, *peak decoding* - defined as the maximal positive peak in a given time window and *peak decoding latency* – defined as the time-point of the maximal positive peak decoding in a given time-window.

## 3. Results

### 3.1. Face category decoding: upright faces vs. houses

We first investigated developmental changes in the time course and overall neural pattern of stimulus categorisation (upright faces vs. upright houses) using a multivariate pattern analysis (MVPA) in each age group. Decoding accuracy, at the group level, was consistently high for all groups, primarily increasing as a function of participant age (peaking at 84.81% for 6-7 year olds, 85.19% 8-9, 78.45% 10-11 and 90.49% adults in comparison to chance levels at around 50%). We also found that significant levels of decoding were reached earlier in the time course (i.e., post presentation of the stimulus) as participant age increased. Adults demonstrated significant decoding most rapidly at 100ms post stimulus onset, followed by the 10-11 year olds at 120ms, then the 8-9 year olds at 128ms and finally the youngest (6-7 year olds) children at 132ms (see Figure 1, top-row, for the time course of decoding accuracy in each group, time-points of significant decoding are highlighted by colour coded dots).

**Figure 1.**
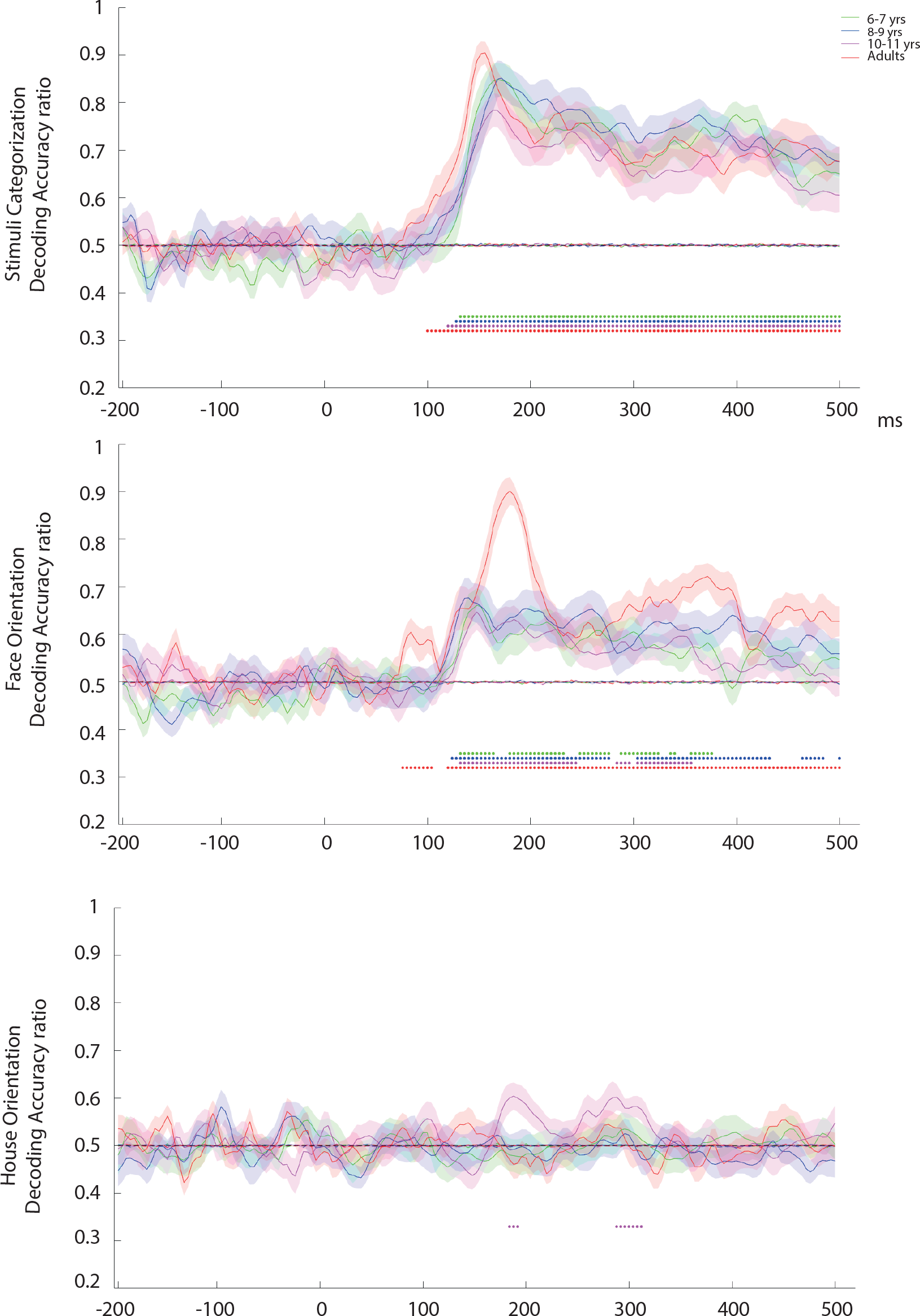
Decoding accuracy comparing upright faces to houses (top row), upright to inverted faces (middle row) and upright to inverted houses (bottom row). Participant age is indicated by colour coding and significant time points are indicated by dots at the base of the curves (p<0.05 (one-tailed) Group-level, FDR corrected).

To formalise these differences and make direct inferential comparisons we extended the standard group analysis by investigating decoding in individual participants. We note that this step is not typically carried out because researchers often rely on group level averages. We confirmed significant decoding in 96% of participants: all but one 6-7 year old and one 10-11 year old (see Supplementary Figure 1 for all individual decoding plots). A between subjects ANOVA (with 4 levels corresponding to the participant age groupings) found no significant effect of participant age on the onset of decoding (F(3,58)= 0.66, p=0.58, η_p_^2^ =0.033^6^), nor on the sustainability of decoding across the epoch (from 60 to 500 ms, F<1). We further compared peak decoding accuracy measured in the time between 100-300 ms (a wide window surrounding the initial main decoding peak identified in all groups at the group level) which did not reveal any significant effects of age group on either the magnitude (F(3,60)= 2.04, p=0.12, η_p_^2^ =0.09) or the latency (F(3,60)= 0.42, p=0.74, η_p_^2^ =0.02) of this peak, see Figure 2 for a visual depiction of these metrics in each age group for category decoding (note the violin plots illustrate individual data points with filled circles, the median of each data set with white circles and the shape of the kernel density estimation of the underlying data distribution in the envelope).

**Figure 2.**
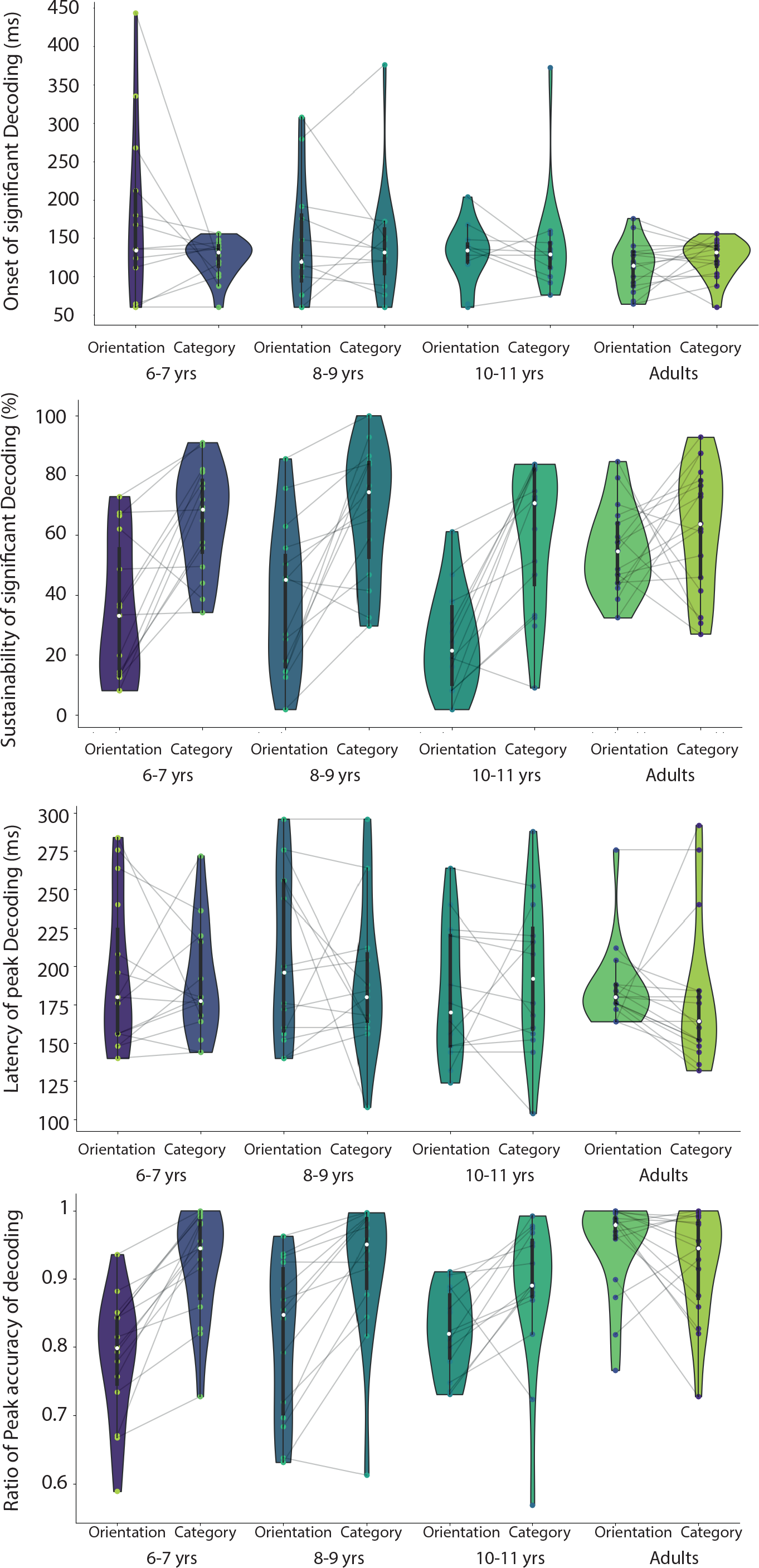
Decoding metrics displayed via violin plots covering onset of decoding (top-row), sustainability of decoding (second row), latency of peak decoding (third row) and amplitude of peak decoding (bottom row). Violin plots highlight the spread of the kernel density estimation of the underlying data distribution (via their envelope), the median of the data set (white dots) and the individual data points. Where possible (i.e. significant decoding was found under both comparisons), for completeness, straight lines link performance for the same individual during Category (upright faces vs. upright houses) and Orientation (upright faces vs. inverted faces) classification.

### 3.2. Face orientation decoding: upright faces vs. inverted faces

We then applied MVPA to investigate the orientation selectivity of decoding for upright vs. inverted faces, as well as our comparison category of houses. At the group level we observed sustained significant decoding of upright vs. inverted face stimuli in all age groups but at much reduced levels in all child groups (accuracy peaked at 66.16% for 6-7 year olds, 67.73% 8-9, 64.78% 10-11 compared with 90.15% in adults; chance levels are around 50%). Furthermore, we again observed that at the group level, significant decoding was reached slightly earlier for the adults at 120 ms (after an initial bump at 76ms), followed by the child groups closely together in time: 6-7 year olds at 132ms; 8-9, 124ms; 10-11, 132ms, see Figure 1, middle-row. Crucially, this sensitivity for stimulus orientation was face selective with no significant decoding of upright versus inverted houses observed in adults, or the youngest child groups (6-7, 8-9 years of age, see Figure 1, bottom-row). The only significant classification of house orientation occurred in two very short time windows in the 10-11 year old children between 184-192ms and 288-31ms.

As before, we extended the analyses to the individual participant level to statistically compare group differences in the onset of significant decoding, the sustainability of decoding, the peak decoding level and the latency associated with the latter. Once again significant face orientation decoding was observed in the majority (92%) of participants (all but two 6-7 year olds and three 10-11 year olds), see Supplementary Figure 2 for all individual classification plots. We observed no significant group difference in the onset of decoding (F(3,50)= 2.07, p=0.116, η_p_^2^ =0.111).^7^ The age-groups differed, however, in the sustainability of decoding over the duration of the epoch (60 to 500ms, F(3,57)= 6.13, p=0.001, η_p_^2^ =0.244). Adults demonstrated a pattern of more sustained decoding (M=55.22±3.48%) relative to children (6-7, M=34.2±6.17%, t(22.35^8^)=−2.97, p=0.07, d= −1.05; 8-9, M=38.68±6.43%, t(21.77)=−2.26, p=0.03, d= −0.80; 10-11, M=24.77±4.51%, t(29)=−5.43, p<0.001, d= −1.96). Furthermore, there was a similar trend for significantly greater decoding in the 8-9yr olds in comparison to the 10-11 year olds (t(27)=−1.75, p=0.092, d= 0.65), but not the 6-7yr olds (t(28)=−0.50, p=0.622, d= −0.18), or between the youngest and oldest children (t(27)=1.22, p=0.23, d= 0.45).

Considering a broad time window around the principal decoding peak for each age group (100ms-300ms), there was a significant effect of participant age group on peak decoding accuracy (F(3,57)=10.71, p<0.001, η_p_^2^ =0.36) which was driven by superior decoding accuracy in adults (M=95.0±1.68%) relative to all child groups, 6-7 yo (M=78.69±2.35%, t(30)=−5.74, p<0.001, d=−2.03), 8-9 yrs (M=82.29±3.10%; t(21.82)=−3.61, p=0.002, d= −1.28) and 10-11 yo (M=82.41±1.64%; t(29)=−5.30, p<0.001, d=−1.91). No significant differences were observed across the child groups (p>0.21). Investigation of the latency of this peak decoding accuracy did not reveal any age-related differences (F(3,57)=0.87, p=0.46, η_p_^2^ =0.04). See Figure 2 for a depiction of these metrics, again shown as violin plots under the heading Orientation. Note that where possible, straight lines connect the equivalent metric for the same individual across the two categorization conditions.

### 3.3. ERP Results

For the standard ERP analysis, we considered the P100 component, both amplitude and latency, and the N170 component amplitude. We used a four-way mixed design ANOVA to investigate the effects of participant age group (6-7, 8-9, 10-11, adults), stimulus category (face, house), stimulus orientation (upright, inverted) and cortical hemisphere (left, right). We focus here solely on the contrasts of direct relevance, i.e., those predicted a-priori from extant literature (a full description of the ERP results can be found in the Supplementary Materials). To this end we report main effects of stimulus category (faces vs. houses) and interactions of category with orientation (upright vs. inverted), and any significant interaction of these factors with participant age group. See Figure 3, top-panel, for the grand-average ERP plots per participant age group, split by experimental stimulus category and cortical hemisphere. Figure 3, lower-panels, depict violin plots illustrating the individual participant statistics for the critical components and experimental conditions (upright and inverted faces, faces and houses) with straight lines connecting participants to visualise the consistency of any difference at the individual participant level.

**Figure 3.**
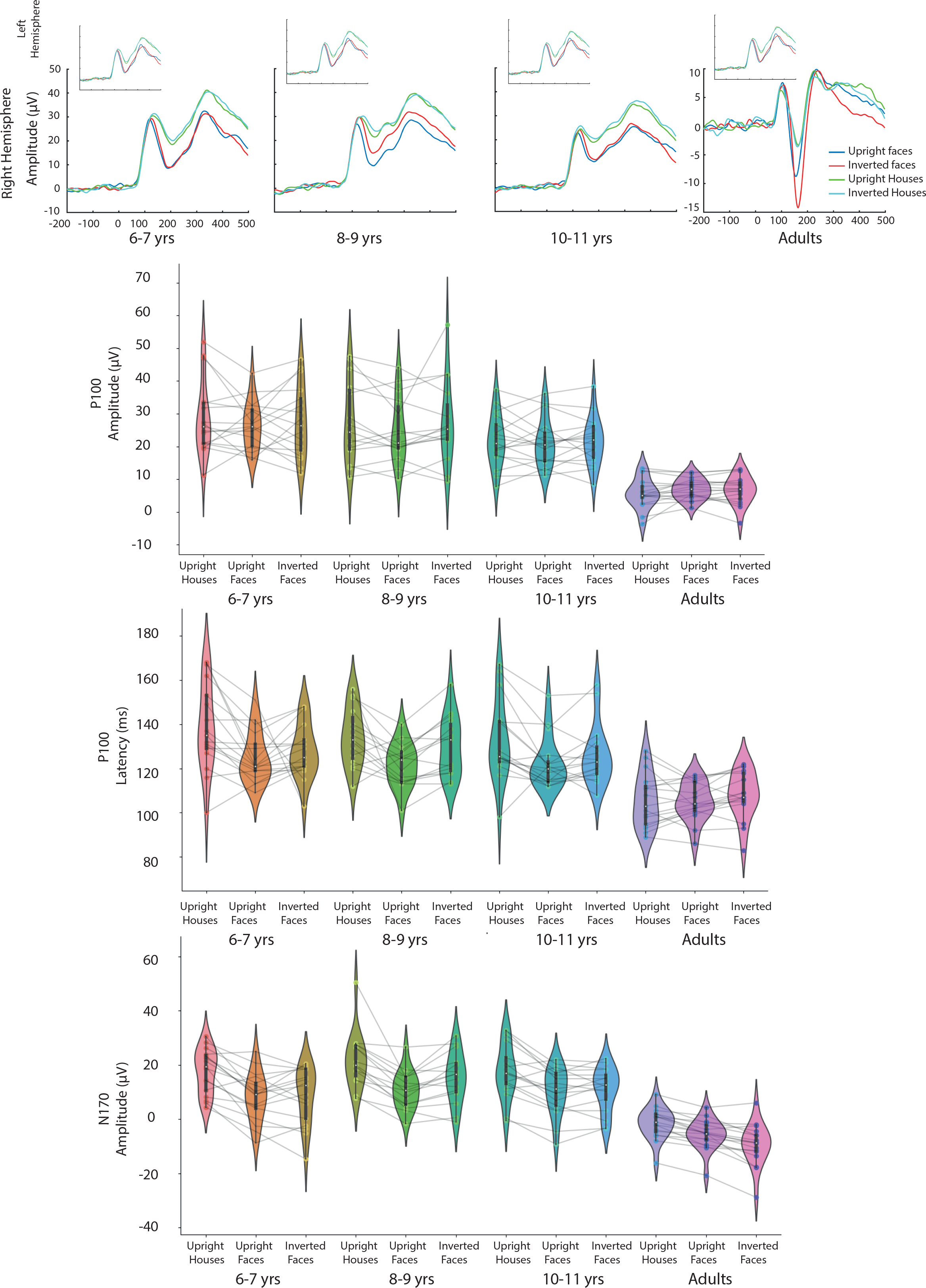
ERP time course for selected right hemisphere electrode (top row, main panel) and left (top row minor panel) for each participant age group. Violin plots (second, third and bottom row) depict the individual data underlying comparisons of the P100 amplitude and latency, and the N170 amplitude in the three critical categories (houses, upright and inverted faces). Straight lines connect the individual data of each participant allowing a direct visualization of the extent to which group level effects are observed on the individual level.

Analysis of **P100 amplitude** revealed a main effect of stimulus category (faces vs. houses) (F(1,62)=4.25; p=0.043, η_p_^2^ =0.06) which reflected a smaller P100 component for faces (M=20.19±1.37 μV) compared to houses (M=21.07±1.58 μV). This effect did not differ as a function of participant age (F(1,62)=1.99; p=0.13, η_p_^2^ =0.09) nor interact further with any other factor or combination of factors (for an interaction between hemisphere and category, F=3.67, p=0.06; F<2.21, p>0.142). There was also a main effect of stimulus category on the **P100 latency** (F(1,61)=37.89; p<0.001, η_p_^2^ =0.38)^9^ which interacted further with participant age-group (F(3,61)=6.60; p=0.001, η_p_^2^ =0.25), and reflected an earlier P100 for faces compared to houses in all child groups (6-7: t(16)=−3.92, p=0.001, d= −0.87; 8-9: t(14)=−4.17, p=0.001, d= −0.81; 10-11: t(15)=−3.75, p=0.002, d = −0.65), but not the adults (t(16)=1.02, p=0.325, d = 0.13). In line with the P100 amplitude there were no further significant interactions of relevance (F<1.44, p>0.24).

A main effect of stimulus category in **N170 amplitude** (F(1,62)=167.97; p<0.001, η_p_^2^ =0.73) reflected a larger response to faces (M=5.96±1.39 μV) than houses (M=14.19±1.61 μV) overall. There was a non-significant trend for this effect to be mediated both by participant age-group (F(3,62)=2.49; p=0.07, η_p_^2^ =0.11)^10^ and by participant age group and stimulus orientation (F(3,62)=2.08; p=0.11, η_p_^2^ =0.91). Probing this latter interaction further to permit clear comparison with the MVPA analysis, significant differences were observed between upright and inverted faces only for the 8-9 year olds (t(14)=−3.39, p=0.04, d= −0.50) and adults (t(16)=4.40, p<0.001, d=0.49), albeit with a reversed profile for the child group (8-9yrs: upright faces: M=10.75±1.83 μV; inverted faces: M=15.30±2.42 μV; adults: upright faces: M=−5.57±1.35 μV; inverted faces: M=−9.46±1.86 μV). No significant differences in N170 amplitude for upright vs. inverted faces were observed for 6-7 or 10-11 year olds (ts<−0.84, ps>0.42, ds<0.09). No differences between upright vs. inverted houses were found for any age group (all ts <−1.41; ps >0.18; ds <0.13).

### 3.4. Summary

On the group level, the MVPA approach indicates significant decoding of both faces versus another object category (houses) and upright versus inverted faces in all age groups tested. Furthermore, this decoding was identified at the individual level in all but a handful of participants. We found no robust evidence for any difference in the latency, sustainability or peak *decoding of face category* decoding (faces vs. houses) as a function of developmental age. In line with this, we found little to distinguish this contrast in children aged 6 – 11 years either from each other, or from adults in the standard ERP analysis, beyond an earlier response to faces in children than adults at the level of the P100 (children, mean between 124.31-126.47 ms, adults M=106.62 ms). There were, however, very clear age-related differences in the more specialised *decoding of face-orientation* in the MVPA approach (NB the same pattern was present in the ERP analysis). The MVPA results indicated that although the distinction between upright and inverted faces can be decoded from the neural response of all of the child groups, adults significantly show a more robust (as indexed by peak decoding magnitude) and sustained (indexed by decoding sustainability) classification of upright faces, relative to inverted, than children. Alongside this, the N170 ERP component analysis also indicated a differential response to face inversion in children and adults. Where adults show the classic enhanced response to inverted faces, this was either entirely absent (6-7yrs, 10-11yrs) or reversed in polarity (8-9yrs) in children. Interestingly, in the MVPA results there was a suggestion that 8-9 year old children differed from their peers in this comparison (with a trend for more sustained decoding than their older peer group).

## 4. Discussion

Questions regarding an early or late maturation of face-selective processing abilities have historically proven difficult to resolve, with mixed findings from the various behavioural studies to date (e.g., Carey & Diamond, 1977; Carey, Diamond, & Woods, 1980; Crookes & McKone, 2009; Germine, Duchaine, & Nakayama, 2011; Hills & Lewis, 2018; Pellicano & Rhodes, 2003; Susilo, Germine, & Duchaine, 2013). The current study attempted to provide clarity on this issue by testing for the presence of distinct profiles of neural activity when children of different ages (and adults) view unfamiliar faces presented in their canonical upright orientation in contrast to inverted. We were particularly interested to see whether any such profile (once observed) is stable or changes across development, in line with increasing face experience and specialist expertise. Using cutting-edge MVPA techniques to probe the neural signal associated with face sensitive processing we present clear evidence that supports the relatively early development of face-selective expertise alongside distinct differences in the strength and extent of face-orientation decoding in children and adults, suggestive of a degree of maturation of the underlying neural processes with age. While the traditional ERP analysis supported the MVPA face-category decoding findings, there was no clear evidence of a differential response to face inversion for children in the standard analysis. Using MVPA in this context permitted a uniquely broad exploration of face selective neural activity, freed from the typical a-priori constraints of predetermined time windows and selected electrodes that are a common and necessary standard for ERP analysis. A more inclusive approach such as this is important when the location and orientation of the neural sources contributing to this category sensitivity in children is known to be highly variable (Scherf et al., 2007) and has provided novel evidence of robust face-orientation decoding across development.

To investigate the tuning of face processing with age and experience, and to probe a hallmark of sophisticated face processing, we contrasted the neural activity associated with upright compared to inverted faces in each of our participant groups. Critically, the novel MVPA analysis of neural activity associated with viewing upright vs. inverted faces indicated that children as young as six have distinct neural representations for upright and inverted faces. This neural face inversion decoding appears to be stable between the ages of 6 – 11 years of age and highly robust as it is observable at the individual level for the majority of participants. Crucially, this differentiation seemed to reflect a particular sensitivity to the canonical upright face orientation, rather than a sensitivity to any change in orientation per se because no such difference was observed for the contrast between upright vs inverted houses. The consistent modulation of neural activity associated with face inversion observed across child age groups converges with evidence of pronounced behavioural effects of face inversion in children (Crookes & McKone, 2009; McKone et al., 2012), which have been observed even in infancy (Turati et al., 2010). Yet our results also reveal that neural differentiation between upright and inverted faces is substantially more pronounced in adults compared to any of the child groups. The relatively greater levels of decoding and more sustained decoding observed in adults supports an ongoing development of face-selective processing abilities between childhood and adulthood. It follows that the adult-like behavioural profile that has been often observed in children may conceal an extended neural maturation of the relevant face networks across development.

Alongside this, the standard ERP analysis suggests that differentiation between upright and inverted faces in the N170 component occurs only for 8-9 yea old children and adults. Moreover, these two groups displayed divergent patterns of activity. As expected, adults showed the typical N170 inversion effect with a higher amplitude for inverted than upright faces (e.g. Bentin, Allison, Puce, Perez, & McCarthy, 1996; Eimer, 2000). By contrast, the 8-9 year olds showed the opposite pattern, with a higher amplitude for upright than inverted faces. Careful interpretation of these results is needed, given the lack of a significant interaction between age group, stimuli category and orientation. Nonetheless, this is not the first observation of a pattern reversal effect for face inversion in children. Indeed, a similar profile was reported previously in a re-analysis combining four separate data sets (see Taylor, Batty, & Itier, 2004) where younger children (8-9yrs) displayed the same pattern reported here but older children (12-15yrs) showed a more adult like pattern. The switch was reported to occur in the 10-11years age bracket where they also observed no difference in N170 response as a function of face inversion. This ‘flipped’ ERP profile, alongside the absence of any significant face inversion effects in the 6-7 and 10-11 year olds, is therefore suggestive of a maturation of face processing networks during childhood, which might be difficult to capture with standard ERP analysis given the high variability in the locus of face-selective areas in children. Such changes are consistent with the fine tuning of face ability with experience claimed by proponents of a late maturation of face specific abilities (Carey & Diamond, 1977; Carey et al., 1980; Germine et al., 2011; Hills & Lewis, 2018; Susilo et al., 2013). In line with these results, several behavioural studies have also noted developmental changes in the face inversion effect (Carey & Diamond, 1977; Hills & Lewis, 2018; Schwarzer, 2000). Similarly, our MVPA results signal that some aspect of fine-tuning of face-inversion representation occurs outside the developmental window examined here, i.e. during late childhood and adolescence.

From a methodological standpoint, the novel application of MVPA approaches presented here yielded insights that would otherwise remain unknown. In particular, we observed very clear and robust evidence of neural differences in the response to face orientation (upright vs. inverted faces) that was entirely absent in the standard ERP responses in two of the age groups tested. The absence of such effects in children aged 10-11 from standard ERP analysis is consistent with previous findings (Taylor et al., 2004a). However, it is now clear that one should not conclude that the absence of such an ERP effect in one analysis approach indicates no difference in the neural response. It is also important to note that the pattern of discriminability is lost in the MVPA analysis e.g., the flipping of the N170 amplitude response as a function of participant age. We would therefore advocate for both approaches as complementary tools towards better characterisation of the underlying neural response profile.

We also compared the neural responses to upright faces and houses to investigate whether children of different ages demonstrate the same basic category sensitivity as adults. We identified distinct face vs. house decoding profiles from around 135ms after stimulus presentation in all age groups overall, and importantly in almost every individual tested. This result provides evidence for an early neural face category sensitivity from 6 years of age, consistent with a hypothesis of early maturation of this face-category distinction. Furthermore, the classic N170 ERP component analysis in the current study also suggests that category sensitivity is relatively stable across the age groups tested, with no evidence of significant change in this effect with developmental age.

An early neural sensitivity to faces as a category of stimuli (compared with other objects) is consistent with the findings of the few ERP studies to have previously targeted this contrast in children (Kuefner et al., 2010; Shen, Lin, Wu, & Chen, 2017; Taylor et al., 2001). In perhaps the most comprehensive investigation to date, Kuefner et al., (2010) analysed N170 sensitivity to faces compared to cars in children and adolescents aged 5 to 16 years and observed no face selective changes across development. Similarly, fMRI investigations find face-preferential activity in children as young as 5 years, albeit with a larger variability in the loci of face sensitivity (Gathers, Bhatt, Corbly, Farley, & Joseph, 2004; Scherf, Behrmann, Humphreys, & Luna, 2007).

Going forward, directly associating developmental changes in brain activity with performance in face related tasks should prove highly informative in understanding the functional impact of the differentiated patterns of neural activation observed here. In particular while there is no question that the face inversion effect reflects something unique about our specialist processing for faces compared to other objects (e.g., Eimer, 2000; Yovel & Kanwisher, 2005), the extent to which face inversion effects can be directly interpreted as an index of configural or holistic processing of upright faces remains unclear (McKone & Yovel, 2009). Tracking changes in these constructs alongside the developmental changes in face related neural activity identified here should deepen our understanding of the maturation of face sensitivity and expertise.

Here we set out to apply state of the art methodological tools to robustly characterise the early neural responses of children aged 6-11 years of age and adults to faces alongside critical comparison categories (objects, inverted faces). Our goal was to bring new evidence to the debate surrounding the typical development of face-processing (broadly contrasting hypothesis of early vs. late maturation of these brain processes). To this end, we provide new findings that both support existing theories and add further complexity to the debate. Our analyses of the EEG response reveal robust profiles of significantly differentiated neural activation associated with viewing faces broadly, i.e., when compared with another stimulus category (houses) and more specifically, i.e., when compared with a stimulus category matched exactly for low level perceptual properties but presented in a non-canonical orientation (inverted faces) from the youngest ages tested. This is indicative of early functional maturation of broad face processing mechanisms. Alongside this we present evidence of ongoing development with age in the form of significant differences in the extent and timing of orientation decoding. Given these findings, it is unsurprising that behavioural studies have reported both impressively expert early face abilities, alongside observations of improvements over time. We hope that future attempts to identify and disentangle the various mechanisms that underpin this developing expertise will benefit from in depth consideration of both neural and behavioural indices, ideally concurrently.

## Acknowledgments

This research was supported by Leverhulme Trust grants RPG-2016-021, RPG-2013-019 awarded to MLS, EF, LE and Annette Karmiloff-Smith. We thank Annette for her invaluable contribution to this project. Her work was an inspiration to us all. We further thank all the children and families who generously gave their time to participate in this project. Many thanks go to Michael Papasavva and Susan Scrimegeour for their help and support during child testing, and to Erin Bartlett for her help with adult testing.

## Footnotes

1 To equate the number of participants in each group we removed 4 participants from the 10-11yrs group and 2 adult participants in the reverse chronological order of testing.

2 11 participants aged 6-7 yrs; 9 participants aged 8-9 yrs; 2 participants aged 10-11 yrs individuals stopped early. No adult participants stopped early.

3 Note that for technical reasons a very small number of participants did not take part in the familiarization task during the EEG set up. An identical pattern of results is observed for the MVPA analysis when these participants are excluded, with the exception of a trend for significantly more sustained face orientation decoding in the 8-9yr olds in comparison to the 10-11 year olds (p=0.092), which is no longer present (p=0.33).

4 Our analyses required approximately equated trial numbers across ages, so we worked to match each group’s mean with the cohort with fewest trials: 6-7 year olds. To this end, we deducted trials from each participant with a surplus (working backwards from the end of their testing session) according to the following formula: 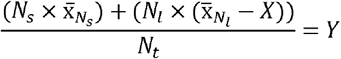 **Y** – Mean number of trials for the target group (in this case, 6-7 year olds); **N**_**S**_ –Number of participants with fewer trials than **Y**; **N**_**l**_–Number of participants with more trials than **Y**; **N**_**t**_ – total number of participants in the age group. Solving this equation allowed us to calculate **X** for each group, which could be removed from each participant with more trials than **Y** to equate the mean number of trials.

5 N.B. To limit the number of multiple comparisons, this analysis was only conducted for time samples between 60 and 500 ms (111 comparisons).

6 Note that two children aged 10-11 did not meet the criterion to establish onset latency and were removed from this analysis.

7 Note that a further two 6-7 yrs; three 8-9 yrs; and two 10-11 yrs participants were removed.

8 Corrected for unequal variance between groups

9 One participant was excluded from this analysis since a peak was not observed in every condition.

10 This trend was driven by a larger difference between N170 amplitude for faces and houses for 6-7 yrs and 8-9 years old children compared to adults (p=0.009; p=0.052; differences between other age groups p>0.14).

