## Supplementary Material for "Developmental changes in the processing of faces as revealed by EEG decoding"

ERP analysis

We ran a mixed ANOVA with repeated factors of stimulus category (faces vs. houses), stimulus orientation (upright vs. inverted), cortical measurement hemisphere (left vs. right) and between subject’s factor of participant age group(6-7, 8-9, 10-11yrs and adults). Results are reported for the P100 amplitude, P100 latency and N170 amplitude in turn. Violations of sphericity are corrected via Greenhouse-Geiser.

**P100 amplitude**

There was a main effect of **stimulus category** (F(1,62)=4.25; p=0.043, η_p_^2^=0.06) driven by a smaller response to faces (M=20.19±1.37 µV) than houses (M=21.07±1.58 µV), a main effect of participant **age group** (F(1,62)=24.13; p<0.001, η_p_^2^=0.54) with a smaller P100 for adults (M=6.40±0.95 µV) compared with all child age groups (6-7 yo: M=27.91±2.17 µV; 8-9: M=26.80±2.93 µV; 10-11: M=22.14±1.83 µV; t>6.63, p<0.001), as well as marginally a smaller P100 for the 10-11 year old children compared to the 6-7 year old children (t(32)=2.03, p=0.051), and a main effect of **hemisphere** (F(1,62)=7.38; p=0.009, η_p_^2^=0.11) with larger amplitudes for the right (M=21.64±1.57 µV) compared to the left hemisphere (M=19.62±1.45 µV). This latter effect of hemisphere further interacted with stimulus orientation (F(1,62)=7.40; p=0.008, η_p_^2^=0.11), such that the effect of orientation was only observed on the right hemisphere (t(65)=-2.78, p=0.007, Cohen’s d=-0.08) with larger amplitudes for inverted stimuli (M=22.21±1.65 µV) compared to upright (M=21.07±1.51 µV), while no difference was observed in the left hemisphere (t(65)=-0.82, p=0.417, Cohen’s d=-0.03; upright: M=19.43±1.41 µV; inverted: M=19.82±1.52 µV). Note that we observed no indication of a significant interaction in the condition of interest, i.e. between stimulus category, orientation and age group: (F(3,62)=0.79; p=0.50, η_p_^2^=0.04) nor between them and hemisphere (F(3,62)=1.64; p=0.190, η_p_^2^=0.07). All other effects of interactions were non-significant (F<3.67, p>0.06).

**P100 latency**

There was a main effect of **stimulus category** (F(1,61)=37.89; p<0.001, η_p_^2^=0.38) driven by an earlier P100 for faces (M=120.62±1.62 ms) compared to houses (M=128.42±2.49 ms), a main effect of **orientation** (F(1,61)=8.06; p=0.006, η_p_^2^=0.12) driven by earlier latencies for upright (M=123.02±1.96 ms) compared to inverted stimuli (M=126.02±2.13 ms), and finally a main effect of participant **age group** (F(3,61)=20.45; p<0.001, η_p_^2^=0.50) driven by an earlier P100 for adults (M=105.88±2.45 ms) compared to all age groups (p<0.001; 6-7 yo: M=133.12±2.80 ms; 8-9: M=131.08±2.97 ms; 10-11: M=129.05±3.21 ms). No differences in latency were found between children (p>0.345). The main effects of **stimulus category and age group** were however mediated by a significant interaction (F(3,61)=6.60; p=0.001, η_p_^2^=0.25) where an earlier P100 was observed for faces compared to houses in all children age groups (6-7 yo: t(16)=-3.92, p=0.001, Cohen’s d= -0.87; 8-9yo: t(14)=-4.17, p=0.001, Cohen’s d= -0.81; 10-11yo: t(15)=-3.75, p=0.002, Cohen’s d = -0.65), but not in adults (t(16)=1.02, p=0.325, Cohen’s d = 0.13). There was no further interaction between stimulus category, orientation and age groups (F(3,61)=0.16; p=0.92, η_p_^2^=0.008) nor between them and hemisphere (F(3,61)=1.44; p=0.24, η_p_^2^=0.07). All other effects of interactions were non-significant (F<2.6, p>0.11).

**N170 amplitude**

There was a main effect of **stimulus category** (F(1,62)=167.97; p<0.001, η_p_^2^=0.73), showing a larger N170 for faces (M=5.96±1.39 µV) than houses (M=14.19±1.61 µV) and a main effect of **age group** (F(3,62)=26.09; p<0.001, η_p_^2^=0.56) driven by a larger N170 for adults compared (M=-4.65±1.44 µV) with all other age groups (6-7 yo: M=13.44±2.02 µV; 8-9: M=17.89±2.39 µV; 10-11: M=14.53±2.09 µV; p<0.001). Furthermore there was a significant **hemisphere by stimulus category** interaction (F(1,62)=11.25; p=0.001, η_p_^2^=0.15), where a left lateralized processing of houses was observed (left: M=13.18±1.57 µV; right: M=15.20±1.77 µV; t(65)=-2.11, p=0.039; Cohen’s d = -0.15) in the absence of any lateralization for face processing (left: M=5.79±1.40 µV; right: M=6.13±1.48 µV; t(65)=-0.44, p=0.66; Cohen’s d = -0.03). There was also an orientation by age group interaction (F(3,62)=7.24; p<0.001, η_p_^2^=.26) where only children aged 8-9 yo (t(14)=-2.95, p=0.011, Cohen’s d =-0.252) and adults were sensitive to stimuli orientation (t(16)=3.98, p=0.01, Cohen’s d= 0.326), but younger children aged 6-7 yrs old (t(16)=-0.98, p=0.34, Cohen’s d =-0.101) and older children aged 10-11 yrs old were not (t(16)=-1.66, p=0.12, Cohen’s d =-0.13). Interestingly while children aged 8-9 yrs old showed a larger N170 for upright stimuli (M=16.27±2.05 µV) compared to inverted (M=19.50±2.80 µV) adults showed the opposite pattern (upright: M=-3.57±1.35 µV; inverted: M=-5.74±1.58 µV).

The interaction of most interest here (between stimulus category, orientation and age groups) indicated a non-significant trend for an interaction only (F(3,62)=2.08; p=0.11, η_p_^2^=0.091) and did not interact further with hemisphere (F(3,62)=0.40; p=0.75, η_p_^2^=0.02). Planned comparisons, however, between upright and inverted faces in each age group did indicate differences between the participant groups. A significant difference between upright and inverted faces was observed in 8-9 yo (t(14)=-3.39, p=0.04, Cohen’s d= -0.50) and adults (t(16)=4.40, p<0.001, Cohen’s d=0.49), but with a reversed polarity in the children. No difference in response to upright faces compared to inverted faces was observed for the younger children 6-7 yrs old (t(16)=-0.32, p=0.76, Cohen’s d= -0.05) or the older children group 10-11 yrs old (t(16)=-.84, p=0.42, Cohen’s d= -0.09). No other effects or interactions reached significance (F<3.50, p>0.07).

Figure S1. Individual classifier accuracy comparing upright faces and upright houses. Classifier accuracy is displayed in red and classifier accuracy trained with permuted labels –chance level- is in blue. Significant time points are indicated by dots at the base of the curves (p<0.05 (one-tailed, FDR corrected; shaded area, SEM).

Figure S2. Individual classifier accuracy comparing upright and inverted faces. Classifier accuracy is displayed in red and classifier accuracy trained with permuted labels –chance level- is in blue. Significant time points are indicated by dots at the base of the curves (p<0.05 (one-tailed, FDR corrected; shaded area, SEM).
